# *GIGANTEA* promotes sorghum flowering by stimulating floral activator gene expression

**DOI:** 10.1101/427492

**Authors:** Frank G. Harmon, Junping Chen, Zhanguo Xin

**Author notes:** **Conflicts of Interest:** The authors declare no conflicts of interest.

## Abstract

**Funding:** This work was supported by USDA-ARS CRIS projects 2030-21000-039-00D and 2030-21000-049-00D to F.G.H.

**Abstract:** The C4 grass *Sorghum bicolor* is an important grain and subsistence crop, animal forage, and cellulosic biofuel feedstock that is tolerant of abiotic stresses and marginal soils. Sorghum is short-day flowering, an obstacle for adaptation as a grain crop but a benefit as a biofuel feedstock. To identify genes underlying sorghum photoperiodic flowering behavior this study characterized the *Sbgi-ems1* nonsense mutation in the sorghum *GIGANTEA* (*SbGI*) gene from a sequenced M4 EMS-mutagenized BTx623 population. *Sbgi-ems1* plants had reduced stature and leaf blades exhibiting increased lateral growth combined with reduced proximal-distal growth. Mutant plants flowered later than normal siblings under long-day conditions provided by greenhouse or field. Delayed flowering in *Sbgi-ems1* plants accompanied by an increase in internode number, indicating an extended vegetative growth phase prior to flowering. *Sbgi-ems1* plants had reduced expression of floral activator genes *SbCO* and *SbEhd1* and downstream FT-like florigen genes *SbFT, SbCN8*, and *SbCN12*. Therefore, *SbGI* accelerates flowering by promotion of *SbCO* and *SbEhd1* expression. Circadian clock-associated genes *SbTOC1* and *SbLHY* had disrupted expression in *Sbgi-ems1* plants. This work demonstrates *SbGI* is a key upstream activator in the regulatory networks dictating sorghum flowering time and growth, as well as gene expression regulation within the circadian clock.

**Summary Statement:** Sorghum *GIGANTEA* contributes to flowering time, growth, and the circadian clock with activities opposite to its maize homolog. *GI* occupies a conserved position within regulatory networks but has plastic activity.

## Introduction

Sorghum is a C4 grass native to Africa that is a key grain and subsistence crop, an animal forage, and a promising cellulosic biofuel feedstock. Advantages of sorghum are its high productivity in marginal soils and under arid conditions. Day length is an important signal for triggering flowering in sorghum and as a short-day (SD) plant flowering is promoted when day length falls below a critical threshold (Craufurd et al., 1999; Quinby, 1974). This sensitivity to day length is at once an obstacle for sorghum adaptation as a grain crop and a benefit for its development as a biofuel feedstock (Mullet et al., 2014). To develop sorghums for either purpose, it is important to understand the genes and regulatory pathways that control photoperiodic flowering.

Photoperiodic flowering is the regulation of flowering time according to day length. This behavior is enforced by an integrated set of transcriptional and post-transcriptional signaling networks that not only promote flowering under inductive photoperiods but also repress flowering under noninductive photoperiods. A highly conserved point of integration for photoperiodic flowering signals is the *CONSTANS* (*CO*) - *FLOWERING LOCUS T* (*FT*) regulatory module (Young Hun Song, Shim, Kinmonth-Schultz, & Imaizumi, 2015), named for genes first discovered in *Arabidopsis thaliana. CO* encodes a member of a family of B-box CCT domain transcription factors widely conserved in plants (Griffiths, Dunford, Coupland, & Laurie, 2003; Putterill, Robson, Lee, Simon, & Coupland, 1995). Arabidopsis *FT* encodes a member of the larger plant PEBP-related family conserved throughout flowering plants that contains smaller group of FT-like florigen-related proteins (Danilevskaya, Meng, Hou, Ananiev, & Simmons, 2008; Turck, Fornara, & Coupland, 2008). According to the florigen hypothesis, a mobile signal originating in leaves transmits the flowering signal to the shoot apical meristem to promote flowering (Pennazio, 2004). Leaf expressed FT-like proteins in Arabidopsis, rice, tomato, and cucurbits serve as molecular florigen signals to trigger the shift from vegetative to floral development at the shoot apical meristem (Jaeger & Wigge, 2007; Lifschitz et al., 2006; Lin et al., 2007; Notaguchi et al., 2008; Tamaki, Matsuo, Wong, Yokoi, & Shimamoto, 2007).

The primary role of *CO* is regulation of *FT* expression and whether CO protein activates or represses its *FT* target genes varies among plants. In Arabidopsis, *CO* activates *FT* expression to promote flowering under floral inductive long-day (LD) photoperiods (Samach et al., 2000; Y.H. Song, Smith, To, Millar, & Imaizumi, 2012; Valverde et al., 2004). In contrast, the rice *CO* ortholog *Heading date 1* (*Hd1*) upregulates expression of the *FT* ortholog *Heading date 3a* (*Hd3a*) in floral inductive SD photoperiods and represses it in LD (Kojima et al., 2002; Yano et al., 2000). *Hd1* also represses expression of rice *Early heading date 1* (*OsEhd1*)*. OsEhd1* encodes a B-type response regulator that promotes *Hd3a* expression under SD separate from *Hd1* (Doi et al., 2004; Itoh, Nonoue, Yano, & Izawa, 2010; Zhao et al., 2015). An upstream activator of *OsEhd1* expression is *Early heading date 2* (*Ehd2*) encoding a zinc finger transcription factor, which is an ortholog of the maize floral activator *INDETERMINATE 1* (*ID*) (Matsubara et al., 2008). The maize *id1* mutant is very late flowering (Colasanti, Yuan, & Sundaresan, 1998). The maize florigen-related gene *Zea mays CENTRORADIALIS 8* (*ZCN8*) is a presumed florigen, since silencing *ZCN8* expression delays flowering (Meng, Muszynski, & Danilevskaya, 2011). *CONSTANS OF Zea mays1* (*CONZ1*) is a co-linear ortholog of rice *Hd1* (Miller, Muslin, & Dorweiler, 2008), but genetic and molecular studies have not tested the contribution of *CONZ1* to regulation of *ZCN8*.

Sorghum *CONSTANS* (*SbCO*) acts upstream to promote expression of *SbEhd1* and several florigen-related genes in both LD and SD photoperiods (Yang, Weers, Morishige, & Mullet, 2014). Of the thirteen PEBP-family genes in sorghum, sorghum *CENTRORADIALIS 8* (*SbCN8*) is the co-linear ortholog of maize *ZCN8* and *SbFT* is the co-linear ortholog of rice *Hd3a* (R L Murphy et al., 2011). An additional PEBP-family gene orthologous between maize and sorghum is *SbCN12* (R L Murphy et al., 2011; Yang et al., 2014). Both *SbCN8* and *SbCN12* possess florigen activity when overexpressed in Arabidopsis (Wolabu et al., 2016). Collectively, *SbFT, SbCN8, SbCN12* are regulated by *SbCO* and *SbEhd1* (Rebecca L. Murphy et al., 2014; Yang et al., 2014), consistent with this set of genes acting as the CO-FT module in sorghum.

An additional repressor of flowering upstream of *SbCO* is the sorghum *PSEUDORESPONSE REGULATOR 37* (*SbPRR37*) (R L Murphy et al., 2011). *SbPRR37* encodes a member of a family of transcriptional repressors originally discovered as core circadian clock genes in Arabidopsis (Farre & Liu, 2013), but *SbPRR37* has no contribution to circadian clock function (R L Murphy et al., 2011). Differentially functional *SbPRR37* alleles underlie the flowering time-associated *Maturity* locus *Ma1*, which has the largest impact on sorghum flowering time (Quinby, 1974). Inactive *ma1/Sbprr37* alleles confer early flowering in LD conditions and played an important role in early domestication of sorghum (Quinby, 1967). Under LD conditions, *SbPRR37* inhibits flowering by repressing expression of flowering activators *SbEhd1* and *SbCO* to ultimately suppress expression of florigen-related genes like *SbFT, SbCN8*, and *SbCN12* (R L Murphy et al., 2011).

*GIGANTEA* (*GI*) is a gene identified in early genetic screens for delayed flowering mutants in Arabidopsis (Koornneef, Hanhart, & van der Veen, 1991; Redei, 1962). *GI* participates in flowering time control, the circadian clock, and a wide range of other physiological activities (Mishra & Panigrahi, 2015). In Arabidopsis, *GI* stimulates flowering by promoting *FT* expression in LD through direct transcriptional regulation of *FT* and post-transcriptional inactivation of *CO* repressors (Park et al., 1999; Sawa & Kay, 2011; Sawa, Nusinow, Kay, & Imaizumi, 2007; Suarez-Lopez et al., 2001). Within the circadian clock, GI protein is fundamental to the protein complex that targets the core circadian clock transcriptional repressor *TIMING OF CAB 1* (*TOC1*) for degradation by the ubiquitin-26S proteasome system (Kim et al., 2007; Mas, Kim, Somers, & Kay, 2003). Tight regulation of TOC1 protein activity is integral to a mutual negative regulatory feedback loop between *TOC1* and another core circadian clock gene *LATE ELONGATED HYPOCOTYL* (*LHY*) (Alabadi et al., 2001; Gendron et al., 2012; Huang et al., 2012). Sorghum has homologs of both *TOC1* and *LHY* (R L Murphy et al., 2011).

*GI* is also an important component of photoperiodic flowering time networks in grasses. *gi* mutants in rice and maize alter flowering time behavior. Under greenhouse conditions the rice *osgi-1* mutant allele delays flowering under SD photoperiods, but not in LD conditions, and only slightly delays flowering in the field (Izawa et al., 2011). *OsGI* is important for blue light-promoted induction of rice *Ehd1* expression as part of the mechanism for critical SD day-length recognition (Itoh et al., 2010). Maize has two paralogous *GI* genes, *GIGANTEA1* (*GI1*) and *GIGANTEA2* (Miller 2008; Mendoza 2012). *gi1* mutants flower earlier in LD, but not SD, and have elevated expression of *ZCN8* and *CONZ1*, indicating *GI1* is an upstream repressor in LD (Bendix, Mendoza, Stanley, Meeley, & Harmon, 2013).

The role played by *SbGI* in regulation of sorghum flowering is not well characterized. A comparative genomic study of 219 African sorghum accessions identified single nucleotide polymorphisms (SNPs) at *SbGI* significantly associated with photoperiod sensitivity (Bhosale et al., 2012). Two associated SNPs caused non-synonymous amino acid changes and a third represented a frameshift mutation. *SbGI* expression has a diel rhythm like all known *GI* genes where peak expression occurs 8 to 10 hours after dawn and this timing is independent of photoperiod (R L Murphy et al., 2011). *SbPRR37* does not contribute substantially to regulation of *SbGI* (R L Murphy et al., 2011).

Here we describe a novel mutant allele in the *SbGI* gene, *Sbgi-ems1*, identified in a sequenced M4 EMS-mutagenized population (Jiao et al., 2016). Plants carrying this nonsense mutation, which truncates GI protein by two thirds, exhibited a number of alterations in growth and development compared to non-mutant normal siblings. Mutant plants had reduced stature and changes in the orientation of leaf blade growth. The *Sbgi-ems1* allele delayed flowering under LD photoperiod conditions provided by greenhouse or field. Delayed flowering in *Sbgi-ems1* accompanied an increase in internode number, indicating an extended vegetative growth phase prior to flowering. This sorghum allele also resulted in reduced expression of the floral activators *SbCO* and *SbEhd1*, as well as limited expression of the FT-like florigen genes *SbFT, SbCN8*, and *SbCN12*. These observations indicate *SbGI* promotes *SbCO* and *SbEhd1* expression, which accelerates flowering time under LD photoperiods. Circadian clock gene expression also was disrupted in *Sbgi-ems1* plants. *SbTOC1* expression was elevated and *SbLHY* expression strongly reduced, consistent with *SbGI* playing an important role in the regulatory network of the sorghum circadian clock.

## Materials and Methods

### Plant stocks and environmental conditions

All sorghum lines are the BTx623/ATx623 genetic background. The ARS223 line is from a collection of 256 whole genome sequenced M4 EMS-mutagenized sorghums lines described by Jiao et al. (2016). Plants were screened for the *Sbgi-ems1* mutation in *SbGI* by Derived Cleaved Amplified Polymorphic Sequences PCR with the primers in Supplemental Table S1. The PCR fragment amplified from the *Sbgi-ems1* locus was resistant to the XcmI restriction enzyme (New England Biolabs) and the product from normal *SbGI* locus was cleaved by this enzyme. Screening of 24 plants from the M4 ARS223 population yielded one plant heterozygous for the *Sbgi-ems1* allele and this plant was used as pollen donor for a cross to a male sterile ATx623 panicle. Progeny of this cross were used for all subsequent experiments.

Plants in the greenhouse were grown under LD conditions of 16-hour days and 8-hour nights. Natural sunlight was supplemented with LumiGrow Pro325 LEDs. Daytime temperature was set to 26°C and nighttime temperature was set to 20°C. Seedlings for growth measurements and gene expression were sown in 4-inch peat pots filled with SuperSoil (The Scotts Company), supplemented with a ½ teaspoon of 14-14-14 N-P-K slow release fertilizer. Plants for flowering experiments were started in the same fashion then transplanted when seedlings reached the 3-leaf stage (10 days-old) to 13-liter pots filled with corn soil (composed of aged wood fines, green waste compost, fir bark, grape compost, rice hulls, chicken manure, red lava, and sandy loam mixed by American Soil and Stone, Richmond, CA). Greenhouse plants were watered twice daily and received 20-20-20 N-P-K fertilizer once a week after being transplanted to 13-liter pots. Field grown plants were maintained in rows at Oxford tract on the University of California, Berkeley campus and watered to soil saturation once weekly by drip irrigation. For each trial,field grown plants were started from seed directly at Oxford tract or transplanted as 4-5 leaf individuals (2 weeks-old) started is peat pots as above. Plants in the field were grow at the Oxford tract on the UC Berkeley campus from late May 2018 to September 2018.

### Growth measurements

Plants were grown under greenhouse conditions to the 6-7 leaf stage (4-6 weeks) and measured at this point for height and leaf blade dimensions. Height corresponded to the distance between the soil surface and the collar of the newest fully expanded leaf. The 6^th^ leaf was dissected from the same plants and its length measured from the tip to the ligule. Width was measured at the midpoint of the blade, determined by folding the leaf blade in half. The same measurements were made for the 6^th^ leaf below the flag leaf from post-flowering *Sbgi-ems1* and normal plants. Internodes above the first internode with prop roots were counted on post-flowering plants from flowering time trials.

### Flowering time

Plants grown under greenhouse conditions were individually scored for the number of days from sowing to reach boot stage and flowering, while field grown plants were scored for boot stage only, due inhibition of anthesis and stigma exertion by the cool temperatures at the Oxford tract. Boot stage was scored as the first day when the entire flag leaf collar was visible in the leaf whorl. Flowering stage was scored as the first day of anthesis for fertile plants or stigma exertion for male sterile plants.

### Gene Expression

Greenhouse grown 6^th^ leaf stage plants were sampled at 0, 8 and 16 hours after dawn. Dawn was when supplemental lights came on at 7 AM. Leaf samples were taken by cutting directly across the 6^th^ leaf ligule with scissors. Two biological replicates were collected for each genotype at each time point. A biological replicate consisted of pooled tissue from three individuals of the same genotype. Leaf samples were flash frozen in liquid nitrogen. After tissue was ground under liquid nitrogen, total RNA was extracted with TRIzol Reagent (ThermoFisher Scientific) according to the manufacturer’s recommendations. 1.5 μg to total RNA for each sample was treated with dsDNase (ThermoFisher Scientific) to remove contaminating genomic DNA and used as a template for cDNA synthesis with the Maxima H Minus First Strand cDNA synthesis Kit (ThermoFisher Scientific) according to the manufacturer’s recommendations. cDNA diluted in half with water served as template for two technical replicate real-time quantitative PCR (qPCR) reactions composed and performed as previously described (Bendix et al., 2013). qPCR reactions for normalization employed PCR primers for 18S RNAs (Supplemental Table S1) and cDNA diluted an additional 1:4000 in water. C_q_ values were calculated with the regression function for each primer set in the Bio-Rad CFX Manager Software (BioRad) and relative transcript levels calculated as 2^(*Cq^18S^*-*Cq^experimental^*).

## Results

### *gi-ems1* is a nonsense EMS mutation in sorghum *GI*

A single *GI* gene is present in sorghum genome on the short arm of chromosome 3 (position 3:3,821,973-3,830,666; Sobic.003G040900; SORBI_3003G040900). Publicly available RNA-seq analysis shows *SbGI* is widely expressed in juvenile and adult tissues, with expression higher in leaf, shoot, and root-related tissues compared to flower- and seed-associated tissues (Supplemental Fig. S1A). The sorghum GI protein is over 95% identical to maize orthologs GI1 and GI2 and 68% identical to the Arabidopsis GI protein (Supplemental Dataset S1).

To evaluate the function of *SbGI*, we took advantage of an existing mutant allele in a collection of M4 EMS-mutagenized BTx623 lines described previously (Jiao et al., 2016). The ARS223 line carries an EMS-induced G to A mutation in *SbGI* at position 5,656 (Fig. 1A). This mutant allele, named here *Sbgi-ems1*, introduces a premature stop codon in place of a conserved tryptophan (W463*). This allele truncates the normally 1162 residue SbGI protein by two thirds to a 462 amino acid protein (Fig. 1A; Supplemental Dataset S1). Individual plants carrying the *Sbgi-ems1* allele were identified in the original ARS223 material by PCR genotyping and the nature of the mutation confirmed by sequencing. One carrier of the *Sbgi-ems1* allele was crossed to a male sterile ATx623 individual to complete backcross 1 (BC1). The BC1F2 and BC1F3 generations were used to evaluate the function of *SbGI*. BC1F3 plants homozygous for *Sbgi-ems1* have reduced overall and peak expression of *SbGI* compared to normal siblings (Fig. 1B), as is common for nonsense alleles. Rhythmic *SbGI* expression persists in *Sbgi-ems1* with peak transcript levels occurring 8 hours after dawn similar to normal plants (Fig. 1B). The nature of the *Sbgi-ems1* mutation and the reduction in gene expression indicate this allele causes significant disruption of *SbGI* activity.

**Figure 1.**
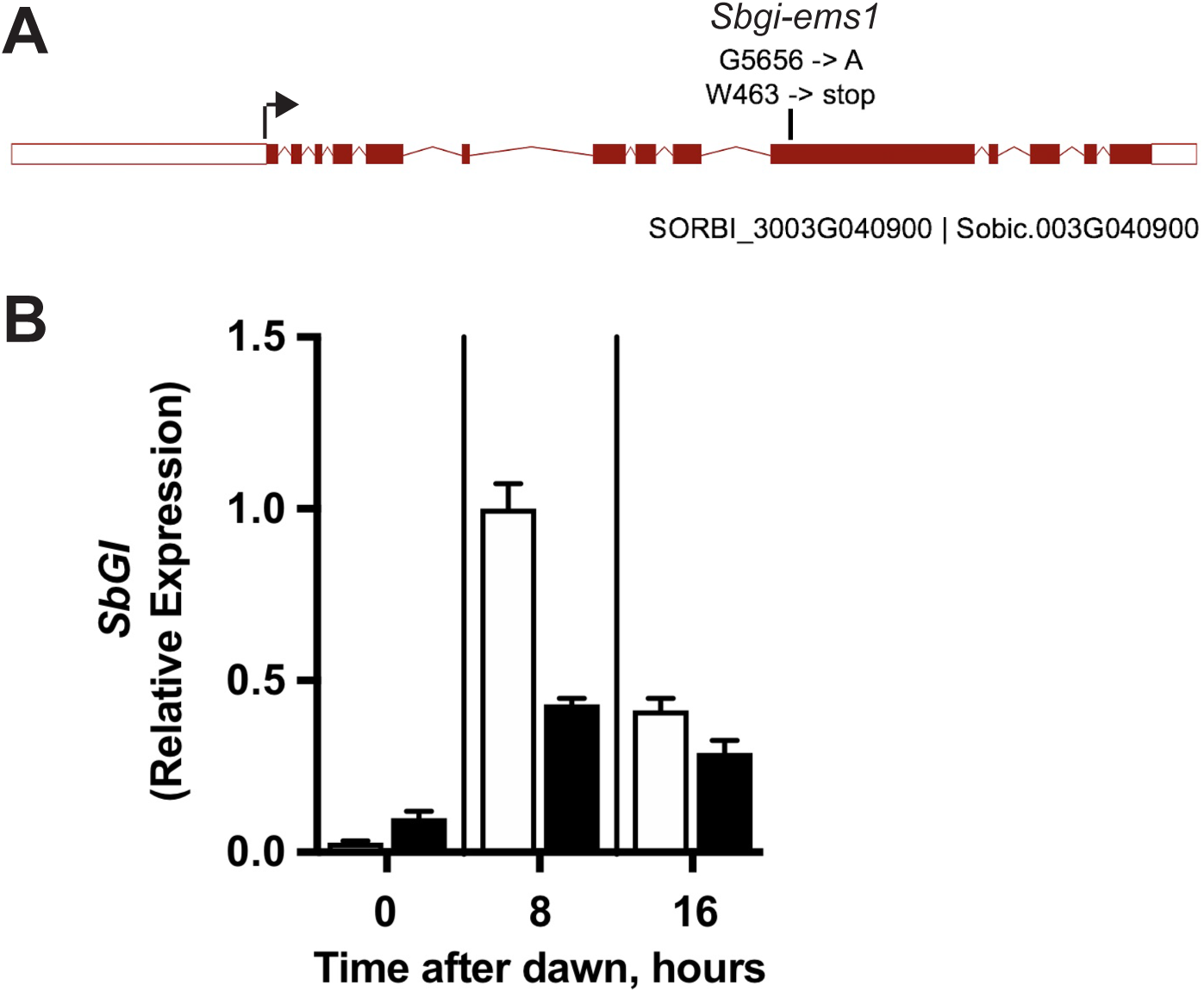
The *Sbgi-ems1* mutant is a nonsense allele that results in reduced *SbGI* expression. A) Diagram of *SbGI* gene (Sobic.003G040900; SORBI_3003G040900) where boxes indicate exons and lines introns. Red coloring indicates coding sequence with the arrow at start codon and white regions indicating 5’- and 3’-UTRs. Vertical line above indicates the position and nature of the *Sbgi-ems1*. B) Transcript levels for SbGI in leaves of normal (white bars) and *Sbgi-ems1* (black bars) BC1F3 plants at 6^th^ leaf stage grown under LD conditions. Time after dawn is the number of hours after lights on in the morning. Values are the average of two biological replicates normalized to the time point from normal plants with highest transcript levels. Error bars represent the range of two biological replicates.

### Visible effects of *Sbgi-ems1* on plant growth

Juvenile *Sbgi-ems1* plants exhibited a clear reduction in stature relative to normal siblings. At the 6-7 leaf stage, homozygous F3 mutant plants were on average 3-5 cm shorter in two trials under LD greenhouse conditions (Fig. 2A). At the same stage, leaf blade growth was also altered in *Sbgi-ems1* plants. Blades from the 6^th^ leaf from juvenile plants were reduced in length and wider at the midpoint (Fig. 2B; Supplemental Fig. S2A, B), leading to a reduction in the length:width ratio in mutant leaves (Fig. 2C). Mature *Sbgi-ems1* individuals at the pre-flowering stage were also visually shorter than normal siblings grown under LD conditions (Fig. 2D). Mature *Sbgi-ems1* plants also exhibited a significant alteration in leaf blade growth, evident as a reduction in the length:width ratio at the midpoint of the 6^th^ leaf below the flag leaf (Fig. 2E). The blade growth change in *Sbgi-ems1* mature leaves is mostly due to an increase in blade width (Supplemental Fig. S2 C, D). These observations indicate *SbGI* activity is important for regulation of both stem and leaf growth in the lateral and proximal-distal directions.

**Figure 2.**
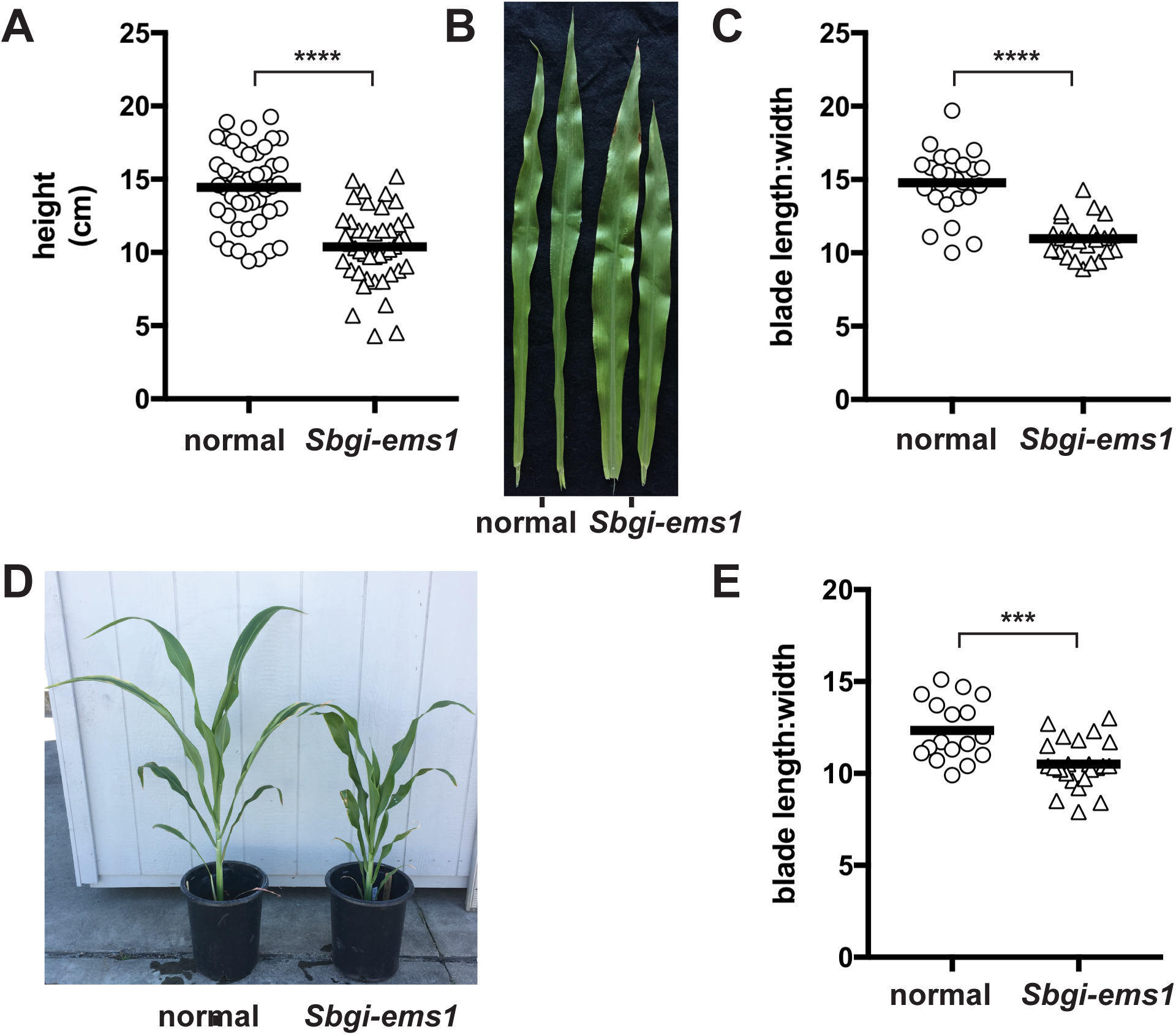
*Sbgi-ems1* reduces plant stature and changes the orientation of leaf growth. Height (A), representative 6^th^ leaves (B), and length:width ratio of blades from 6^th^ leaves (C) of BC1F3 juvenile normal (circles) and *Sbgi-ems1* (triangles) plants grown to the 6–7 leaf stage. D) Representative 2-month-old pre-flowering normal and *Sbgi-ems1* plants. E) Length:width ratio of blades from 6^th^ leaf below flag leaf on BC1F3 mature post-flowering normal (circles) and *Sbgi-ems1* (triangles) plants. All plants were grown under LD greenhouse conditions. Length:width ratio was calculated from length and width measurements in Supplemental Figure S2. All measurements are shown from two separate trials, bar represents the average of measurements. Statistical significance is indicated according to a two-tailed unpaired t-tests with Welch’s correction at P value <0.0001 (****), <0.001 (***), <0.01 (**) and <0.05 (*).

### *Sbgi-ems1* causes delayed flowering

The *Sbgi-ems1* is associated with delayed flowering time under LD greenhouse and field conditions. To assess whether *SbGI* contributes to sorghum flowering time, a total of 114 individuals from the BC1F2 population were scored for flowering time under LD greenhouse conditions in three separate trials. Flowering time was initially scored as days to anthesis for male fertile plants and days to the exertion of stigma for male sterile plants. The timing of each of these events was indistinguishable within the BC1F2 and BC1F3 groups of plants genotyping as normal at *SbGI* (Supplemental Fig. S3A). The group of plants homozygous for the *Sbgi-ems1* allele consistently reached anthesis/stigma exertion an average of 35 days later than *gi1-ems1* heterozygous and normal siblings (Fig. S3A). Heterozygous *Sbgi-ems1* plants reached flowering an average of a week later than normal plants. The late flowering trait tightly co-segregated with the homozygous *Sbgi-ems1* genotype in this BC1F2 population (Supplemental Fig. S3B). Two BC1F3 lines each for *Sbgi-ems1* and normal plants were selected from this BC1F2 population for further analysis.

Delayed flowering time was also evident for *Sbgi-ems1* BC1F3 lines. In two separate greenhouse trials, anthesis or stigma exertion for each BC1F3 *Sbgi-ems1* population occurred an average of 20 days later than the normal F3 lines under LD conditions (Fig. 3B). The timing of boot stage, which occurs prior to anthesis, was determined in these trials to get a more complete idea of the aspect of flowering changed by *Sbgi-ems1*. Similar to the timing of anthesis, boot stage occurred an average of 20 days later in *Sbgi-ems1* plants (Fig. 3B). The timing of boot stage was also determined for the third trial with the BC1F2 population. *Sbgi-ems1* homozygotes in this population were delayed reaching boot stage compared to normal and heterozygous plants (Supplemental Fig. S3C). The average number of days between boot stage and anthesis/stigma exertion for *Sbgi-ems1* and normal plants was not different in all greenhouse trials and in the third trial with the BC1F2 population (Supplemental Fig. S3D). Thus, the flowering time phenotype of *Sbgi-ems1* plants arises from a delay in achieving boot stage, instead of lengthening of the time from boot stage to anthesis.

**Figure 3.**
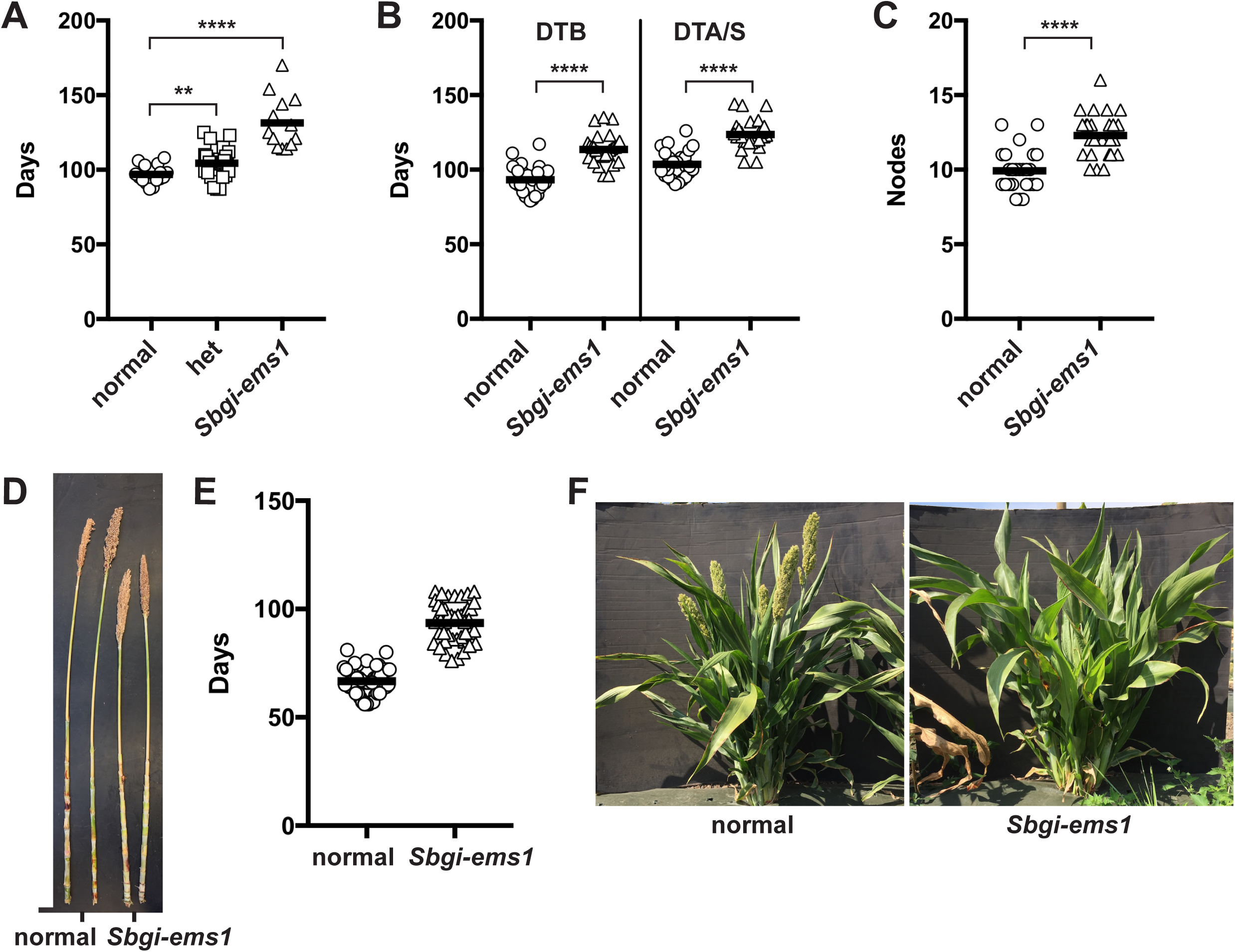
*Sbgi-ems1* mutants are late flowering and produce more internodes prior to flowering. A) Flowering time for BC1F2 population for normal (circles), heterozygous *Sbgi-ems1/SbGI* (squares), and *Sbgi-ems1* (triangles) plants grown under LD greenhouse conditions determined as days to anthesis (fertile plants) or stigma exertion (male sterile plants). B) Flowering time for BC1F3 normal (circles) and *Sbgi-ems1* (triangles) plants grown under LD greenhouse conditions determined as days to boot stage (DTB) and days to anthesis or stigma exertion (DTA/S). C) Number of internodes (Nodes) above prop roots produced by normal (circles) and *Sbgi-ems1* (triangles) BC1F3 plants from flowering time experiments. D) Representative main stems after leaf removal from normal and *Sbgi-ems1* plants from flowering time experiments under greenhouse conditions. E) Flowering time for BC1F3 normal (circles) and *Sbgi-ems1* (triangles) plants grown under summer field conditions determined as days to boot stage. F) Pictures of representative 3-month-old normal and *Sbgi-ems1* plants grown in the field. All measurements are shown from two separate trials, bar represents the average of measurements. Statistical significance is indicated according to a two-tailed unpaired t-tests with Welch’s correction at P value <0.0001 (****), <0.001 (***), <0.01 (**) and <0.05 (*).

BC1F3 generation *Sbgi-ems1* plants produced more internodes than normal siblings. The number of internodes were counted for the F3 plants from the greenhouse flowering trials. *Sbgi-ems1* mutant plants made an average of 2 to 3 internodes than normal F3 plants (Fig. 3C). While *Sbgi-ems1* mutants made additional internodes, the length of the main stem of *Sbgi-ems1* plants remained at or below that attained by normal plants (Fig. 3D). These observations are consistent with an extended vegetative growth phase in *Sbgi-ems1* mutant plants, consistent with the *Sbgi-ems1* allele delaying the timing of the vegetative to floral transition.

Flowering time of BC1F3 *Sbgi-ems1* lines was delayed in field grown plants. To test the importance of GI function to the flowering behavior of field grown plants, days to boot stage was determined for F3 *Sbgi-ems1* and normal plants grown under LD summer field conditions in Berkeley CA. Cool temperatures at this field site precluded reliable scoring of anthesis or stigma exertion in all genotypes. The combined results of two trials indicated that *Sbgi-ems1* plants reached boot stage more than 25 days later than normal plants (Fig. 3E, F). Clearly, *SbGI* contributes to the timing of flowering under field conditions, as well as in the greenhouse.

### *Sbgi-ems1* reduces expression of key flowering time genes

The *Sbgi-ems1* allele causes reduced expression of genes that promote flowering. The effect of the *Sbgi-ems1* allele on expression of flowering-related genes was investigated to understand molecular changes underlying delayed flowering. Levels of transcripts for florigen-related genes *SbCN8, SbCN12*, and *SbFT* were assessed at 0, 8, and 16 hours after dawn in month-old normal and *Sbgi-ems1* plants grown under the same greenhouse conditions as the flowering time experiments. In normal plants, *SbCN8* and *SbFT* transcripts reached peak levels 8 hours after dawn (Fig. 4A, B), which coincides with the time of maximal *SbGI* expression (Fig. 1B). *SbCN12* transcripts, on the other hand, were at similar levels in all three time points (Fig. 4C). *Sbgi-ems1* plants had reduced levels of *SbCN8, SbCN12*, and *SbFT* transcripts at all three time points. The greatest change for *SbCN8* and *SbFT* occurred 8 hours after dawn, where *SbCN8* and *SbFT* achieved levels 3- and 5-fold lower levels than normal, respectively. *SbCN12* levels were reduced by more than 8-fold at each time point. These observations are consistent with *SbGI* serving to promote expression of these three florigen-related genes.

**Figure 4.**
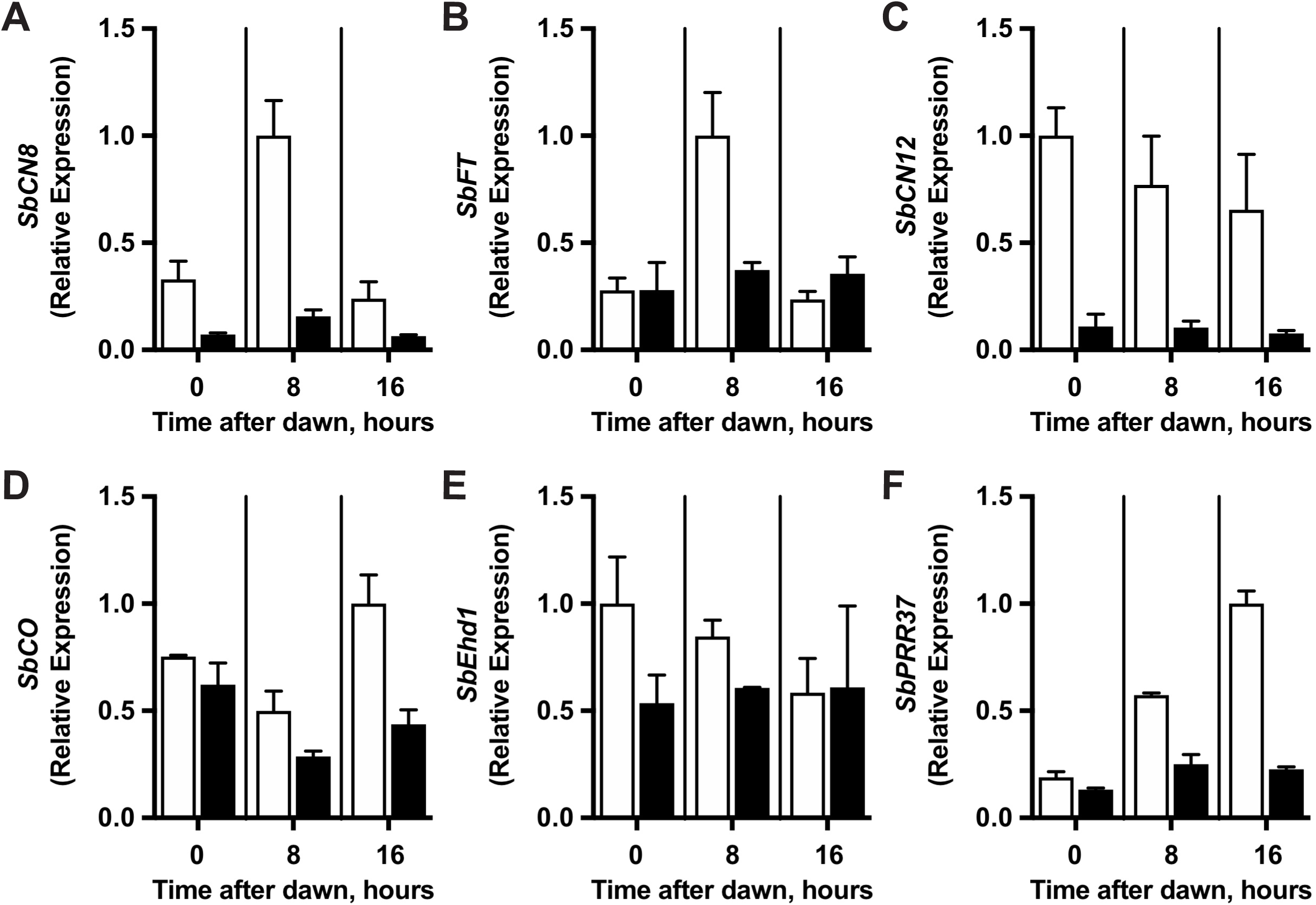
*Sbgi-ems1* alters flowering time gene expression patterns and levels. Transcript levels for *SbCN8* (A), *SbFT* (B), *SbCN12* (C), *SbCO* (D), *SbEhd1* (E), and *SbPRR37* (F) in leaves of normal (white bars) and *Sbgi-ems1* (black bars) BC1F3 plants at 6^th^ leaf stage grown under LD conditions. Time after dawn is the number of hours after lights on in the morning. Values are the average of two biological replicates normalized to the time point from normal plants with highest transcript levels. Error bars represent the range of two biological replicates.

Since the upstream action of *SbCO* and Ehd1 control *SbCN8, SbCN12*, and *SbFT, SbCO* and *SbEhd1* expression was evaluated in normal and *Sbgi-ems1* plants. *SbCO* transcript was present throughout the day in normal plants, with highest levels reached 16 hours after dawn (Fig. 4D). *SbEhd1* expression, on the other hand, was biased toward dawn by 2-fold relative to the 16-hour time point (Fig. 4E). In *Sbgi-ems1* plants, *SbCO* transcript levels were diminished at both 8 and 16 hours after dawn. In normal plants, *SbEhd1* transcript levels were lower in the *Sbgi-ems1* background at all time points and the greatest reduction of 2-fold occurred at dawn. These results indicate *SbGI* promotes expression of the two floral activators *SbCO* and *SbEhd1* under LD conditions.

The expression of the floral repressor *SbPRR37* was also tested to determine whether *SbGI* contributes to its regulation. In normal plants, *SbPRR37* transcript levels peaked 16 hours after dawn (Fig. 4F). At all three time points in *Sbgi-ems1*, the *SbPRR37* transcript was below the basal level observed in normal plants at dawn. Thus, *SbGI* activity contributes to the regulation of *SbPRR37*.

*Sbgi1-ems1* did not alter expression of *SbID1*, a sorghum ortholog of maize ID1, an upstream activator of *SbEhd1* that is not directly regulated by SbCO. In normal plants, the highest levels of *SbID1* transcript occurred at dawn (time 0 hours) and were reduced by half at the time points 8 and 16 hours after dawn (Supplemental Fig. S4A). The *Sbgi-ems1* allele did not change *SbID1* transcript levels at any time point. Therefore, *SbGI* is not involved in the regulation of *SbID1*.

### *gi-ems1* disrupts a core circadian clock feedback loop involving *SbLHY* and *SbTOC1*

The *Sbgi-ems1* allele caused disruption of expression for the circadian clock genes *SbLHY* and *SbTOC1*, which are expected to regulate one another in a negative feedback loop. Since *GI* genes participates in circadian clock function, *Sbgi-ems1* plants were evaluated for changes in expression of the core circadian clock genes *SbTOC1* and *SbLHY. SbTOC1* transcript was evening-expressed with peak levels occurring 16 hours after dawn in normal greenhouse grown normal plants (Fig. 5B), while *SbLHY* transcript was morning-expressed with peak levels occurring at dawn (Fig. 5A). In *Sbgi-ems1* plants, *SbTOC1* transcript was elevated relative to normal levels at all time points, particularly at dawn. On the other hand, the *SbLHY* transcript was not detectable in mutant plants. These observations indicate *SbGI* activity is for needed for proper function of an *SbLHY* and *SbTOC1* regulatory negative feedback loop within the sorghum circadian clock.

**Figure 5.**
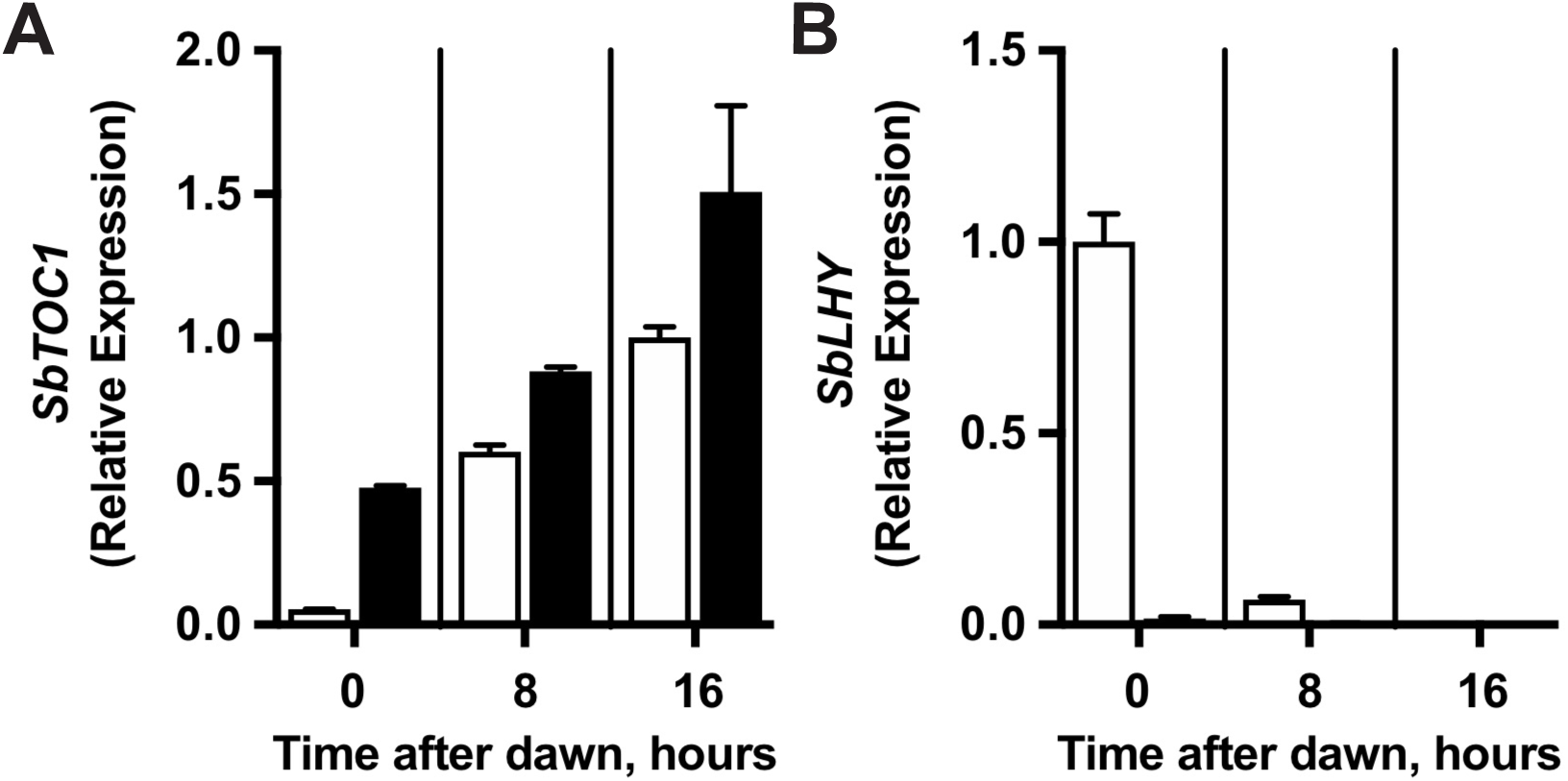
*Sbgi-ems1* disrupts *SbLHY* and *SbTOC1* expression. Transcript levels for *SbLHY* (A) and *SbTOC1* (B) in leaves of normal (white bars) and *Sbgi-ems1* (black bars) BC1F3 plants at 6^th^ leaf stage grown under LD conditions. Time after dawn is the number of hours after supplemental lights came on in the morning. Values are the average of two biological replicates normalized to the time point from normal plants with highest transcript levels. Error bars represent the range of two biological replicates.

## Discussion

Identification of an EMS-derived mutation in the *SbGI* gene, *Sbgi-ems1*, allowed us to evaluate the contribution of *SbGI* to sorghum growth and flowering time. The *Sbgi-ems1* allele is a premature stop codon that truncates GI protein to two thirds of its normal length. Plants homozygous for the *Sbgi-ems1* allele have reduced stature and altered leaf growth. The leaf blade of mutant plants exhibited increased lateral growth and reduced proximal-distal growth, leading to a distortion of the length:width ratio. *Sbgi-ems1* also plants flower later in LD conditions after extended vegetative growth. The delay in flowering is accompanied by a reduction in expression of genes that activate flowering, including the florigen-related genes *SbFT, SbCN8* and *SbCN12*, as well as their upstream regulators *SbCO* and *SbEhd1*. Also, expression of circadian clock genes is disrupted by the *Sbgi-ems1* allele. These observations provide insight into the function of *SbGI*, as well as the regulatory networks that determine flowering time in sorghum.

The flowering behavior of *Sbgi-ems1* mutant plants indicates *SbGI* acts early in control of flowering time. *Sbgi-ems1* delayed flowering time under both greenhouse and field conditions when flowering was scored for BC1F2 and BC1F3 individuals as either days to reach boot stage or days to flowering measured as anthesis (for fertile panicles) or stigma exertion (for sterile panicles); however, the time interval between boot stage and anthesis was unchanged in mutant plants relative to normal or heterozygous plants. Thus, the >20 additional days *Sbgi-ems1* plants required to reach flowering represents a delay in physiological processes leading up to boot stage. These observations indicate the *Sbgi-ems1* allele primarily changes events early in determination of flowering time, not later processes associated with flower development and/or release of pollen/stigma exertion.

An early for role *SbGI* in flowering time is consistent with the observation that *SbGI* is necessary for the proper up-regulation of florigen-related genes *SbFT, SbCN8* and *SbCN12*. In normal plants, *SbFT* and *SbCN8* were rhythmically expressed with peak levels occurring at 8 hours after dawn, while *SbCN12* expression reached similar levels across the day. The midday peak in *SbFT* and *SbCN8* expression coincided with a similarly timed peak in *SbGI* expression. *Sbgi-ems1* plants, on the other hand, had reduced *SbCN12, SbFT* and *SbCN8* expression at each time point. Rhythmic expression of *SbFT* and *SbCN8* was notably absent in *Sbgi-ems1*, instead each transcript was present at a low constant level at each time. These results are consistent with a requirement for *SbGI* activity to promote expression of these three florigen-related genes, in particular midday expression *SbFT* and *SbCN8*.

*SbGI* appears to promote florigen-related expression through up-regulation of *SbCO* and *SbEhd1* expression. *SbCO* and *SbEhd1* stimulate florigen gene expression under both LD and SD conditions (Rebecca L. Murphy et al., 2014; Yang et al., 2014). Additionally, *SbCO* activates *SbEhd1* expression under the all photoperiod conditions. *SbCO* expression is reduced in the *Sbgi-ems1* background, indicating *SbGI* is involved in activation of *SbCO* at the transcriptional level; however, another possibility that cannot be ruled out is SbGI-directed inactivation of an *SbCO* repressor. Also, *SbEhd1* expression is reduced in *Sbgi-ems1*, which could arise from diminished *SbCO* or the absence of direct *SbGI* up-regulation of *SbEhd1*. It is notable that the most significant reduction in *SbEhd1* expression in *Sbgi-ems1* occurs at dawn. This expression pattern is reminiscent of the loss of morning-induced *OsEhd1* expression in the *osgi-1* mutant (Itoh et al., 2010). Thus, *SbGI* may contribute to a regulatory “gate” that promotes *SbEhd1* expression in the morning. *Sbgi-ems1* had no impact on expression of *SbID1*, indicating reduced *SbEhd1* expression in the mutant background in not due to a lack of up-regulation by *SbID1*.

The flowering time delay in *Sbgi-ems1* was unlikely a consequence of reduced SbPRR37 protein-directed repression of *SbEhd1* and *SbCO* even though *SbPRR37* expression was diminished in the mutant background. The BTx623/ATx623 genetic background used here carries the *Sbprr37-3* allele of *ma1* that encodes inactive PRR37 (R L Murphy et al., 2011). Nevertheless, the change in *SbPRR37* expression in *Sbgi-ems1* provides insight into regulation of *SbPRR37*. Lower *SbPRR37* expression in the *Sbgi-ems1* background could arise from either loss of direct activation by *SbGI* or an indirect result of a marked disruption of the circadian clock. In the latter case, strong repression of *SbPRR37* may result from elevated *SbTOC1* expression. In the Arabidopsis clock system, *TOC1* represses *PRR7* as part of a timed series of repressive events involving a suite of PRR-family genes (Pokhilko et al., 2012). A reciprocal effect of *SbPRR37* on *SbTOC1* is unlikely, since *SbPRR37* is not a component of the sorghum circadian clock as shown by the absence of circadian clock defects in *Sbprr37/ma1* alleles (R L Murphy et al., 2011).

Comparing the observations here for *Sbgi-ems1* to previous work on maize *gi1* mutants highlights interesting differences in the roles played by *GI1* and *SbGI* in these related C4 grasses. The sorghum *Sbgi-ems1* allele and maize *gi1* mutants change growth and flowering time in opposite directions. While the sorghum *Sbgi-ems1* mutant caused significantly later flowering under LD conditions, flowering time is modestly accelerated in maize *gi1* mutants under LD photoperiods (Bendix et al., 2013). Additionally, sorghum *Sbgi-ems1* plants had reduced stature, while maize *gi1* mutants grow taller. It is possible to infer from analysis of gene expression that differences in flowering time between maize and sorghum arise from opposite activities for the cognate *GI* gene. *SbGI* serves as an activator of *SbCO*, leading to reduced expression of *SbCO, SbEhd1*, and downstream florigen-related genes *SbFT, SbCN8*, and *SbCN12* in *Sbgi-ems1*, while maize *gi1* is a repressor of *CONZ1*, leading to upregulation of *CONZ1* and *ZCN8* in *gi1* mutant backgrounds.

In both maize or sorghum, the genesis of growth changes observed in *gi* mutants remains unclear. Since Arabidopsis GI protein has been implicated in gibberellin (GA) signaling (Tseng et al., 2004) and *OsGI* is needed for proper regulation of GA biosynthesis genes (Itoh & Izawa, 2011), it is possible that alterations in gibberellin biosynthesis or signaling underlie the growth phenotypes observed in sorghum and maize *gi* mutants. If this is the case, then the sorghum and maize GI proteins are predicted to have opposite regulatory roles there as well.

The *Sbgi-ems1* allele disrupted expression of the circadian clock genes *SbLHY* and *SbTOC1*. This is consistent with alteration of a mutual regulatory feedback loop between *SbLHY* and *SbTOC1* similar to that described in the Arabidopsis circadian clock (Alabadi et al., 2001; Gendron et al., 2012; Huang et al., 2012). Interestingly, *TOC1* expression increases and *LHY* decreases in the *osgi-1* mutant background (Izawa et al., 2011). The similar effect of sorghum and rice *gi* mutants on *SbTOC1* and *SbLHY* expression indicates comparable circadian clock regulatory networks exist in these grasses. Also, the SbGI and OsGI proteins appear to be involved in transcriptional control of *TOC1* expression. By contrast, Arabidopsis GI protein serves to regulate TOC1 protein activity at the post-transcriptional level (Kim et al., 2007; Martin-Tryon, Kreps, & Harmer, 2007). These observations indicate regulation of *TOC1* activity by *GI* is a conserved feature of plant circadian clocks, but the underlying molecular mechanisms are potentially different between plant species.

## Acknowledgements

Thank you to Emma Kovak and Carine Marshall for feedback and advice. We thank Lia Poasa and Julie Calfas at PGEC greenhouse and Tina Wistrom, and Al Hunter at UC Berkeley Greenhouse Facilities for excellent care of plants. Julianne Elliot and Parkesh Suseendran provided invaluable technical assistance. This work was supported by USDA-ARS CRIS projects 2030-21000-039-00D and 2030-21000-049-00D to F.G.H.

## vii. Acknowledgements

Thank you to Emma Kovak and Carine Marshall for feedback and advice. Julianne Elliot and Parkesh Suseendran provided invaluable technical assistance. We thank Lia Poasa and Julie Calfas from the PGEC greenhouse and UC Berkeley Greenhouse Facilities staff Tina Wistrom and Al Hunter for excellent care of plants. This work was supported by USDA-ARS CRIS projects 2030-21000-039-00D and 2030-21000-049-00D to F.G.H.

## Authorship

Xin and Chen generated and characterized the sequenced EMS-mutagenized sorghum population containing the *Sbgi-ems1* allele. Chen performed early experiments on populations with the *Sbgi-ems1* allele. Harmon conceived, designed, and performed the research. Harmon wrote the manuscript with input from the co-authors.

## Supplemental Files

Supplemental Table S1. Primers used in this study.

Supplemental Figure S1. *SbGI* expression in various sorghum tissues.

Supplemental Figure S2. Length and width measurements of leaf blades from juvenile and mature plants.

Supplemental Figure S3. Late flowering of *Sbgi-ems1* plants arises from delayed boot stage.

Supplemental Figure S4. Expression of floral activator *SbID1* in *Sbgi-ems1*.

Supplemental Dataset S1. Amino acid alignment of GI proteins from sorghum, maize, and Arabidopsis.

